## Supplementary Information for "A natural *timeless* polymorphism allowing circadian clock synchronization in ‘white nights’"

Supplementary Figure 1

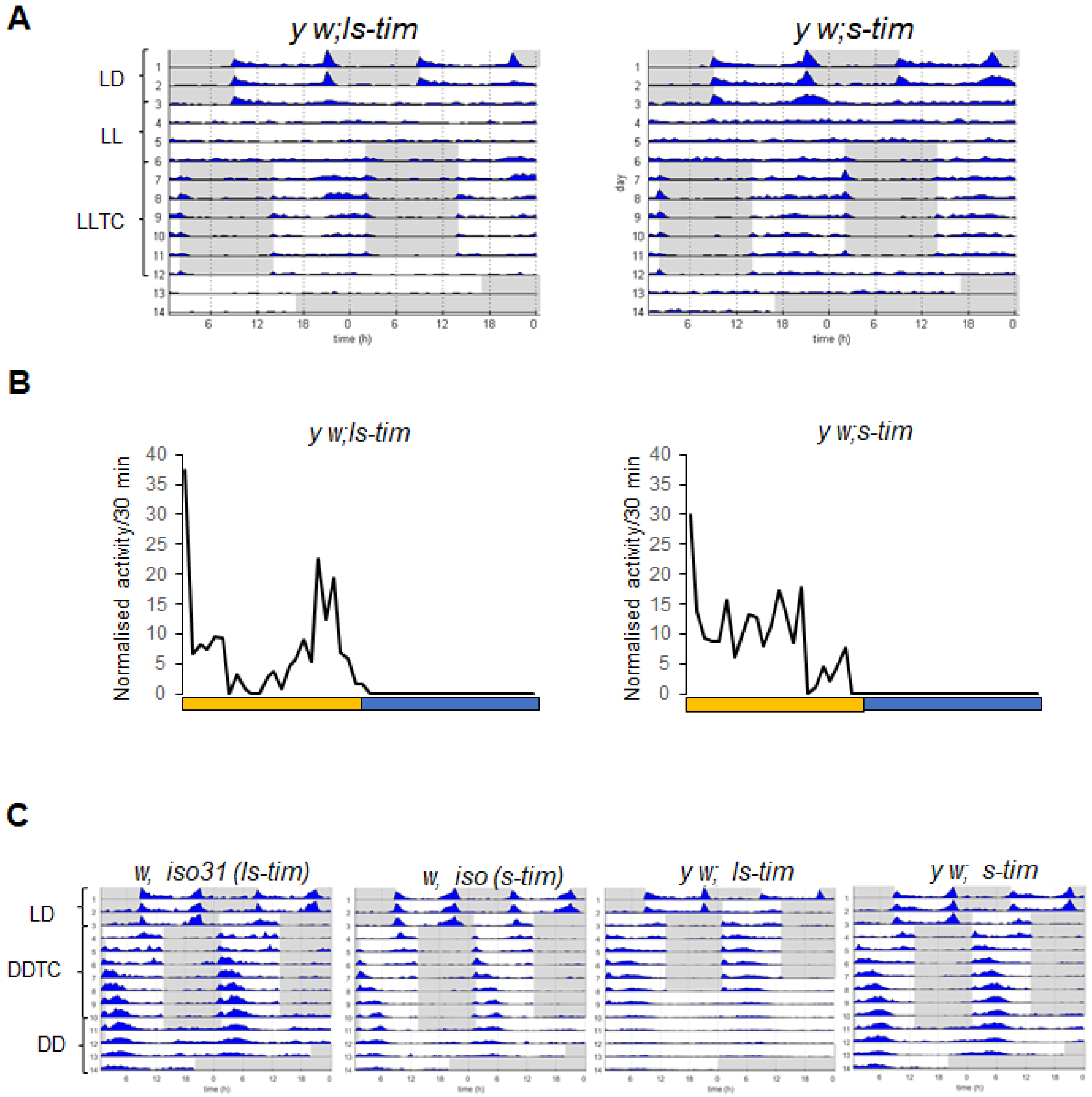


**Supplementary Figure 1: *s-tim* flies cannot synchronize their behaviour to temperature cycles in constant light. A)** Group actograms of one representative experiment as described in the legend for Figure 1A. N (*y w; ls-tim*): 22, (*y w; s-tim*): 18. **B)** Median of normalised activity during day 6 of LLTC of independent experiments combined. Yellow bar: thermophase, blue bar cryophase (12h each). N (*y w; ls-tim*): 50, (*y w;s-tim*): 52. **C)** Group actograms in LD followed by DDTC and DD constant temperature. White areas: lights-on and 25°C during LD, and lights-off and 25°C during DDTC and DD. Grey areas: lights-off and 25°C during LD, and lights-off and 16°C during DDTC. N (*w, iso31 ls-tim*): 15; (*w, iso s-tim*): 15; (*y w; ls-tim*): 15; (*y w; s-tim*): 17.

**Supplementary Figure 2**

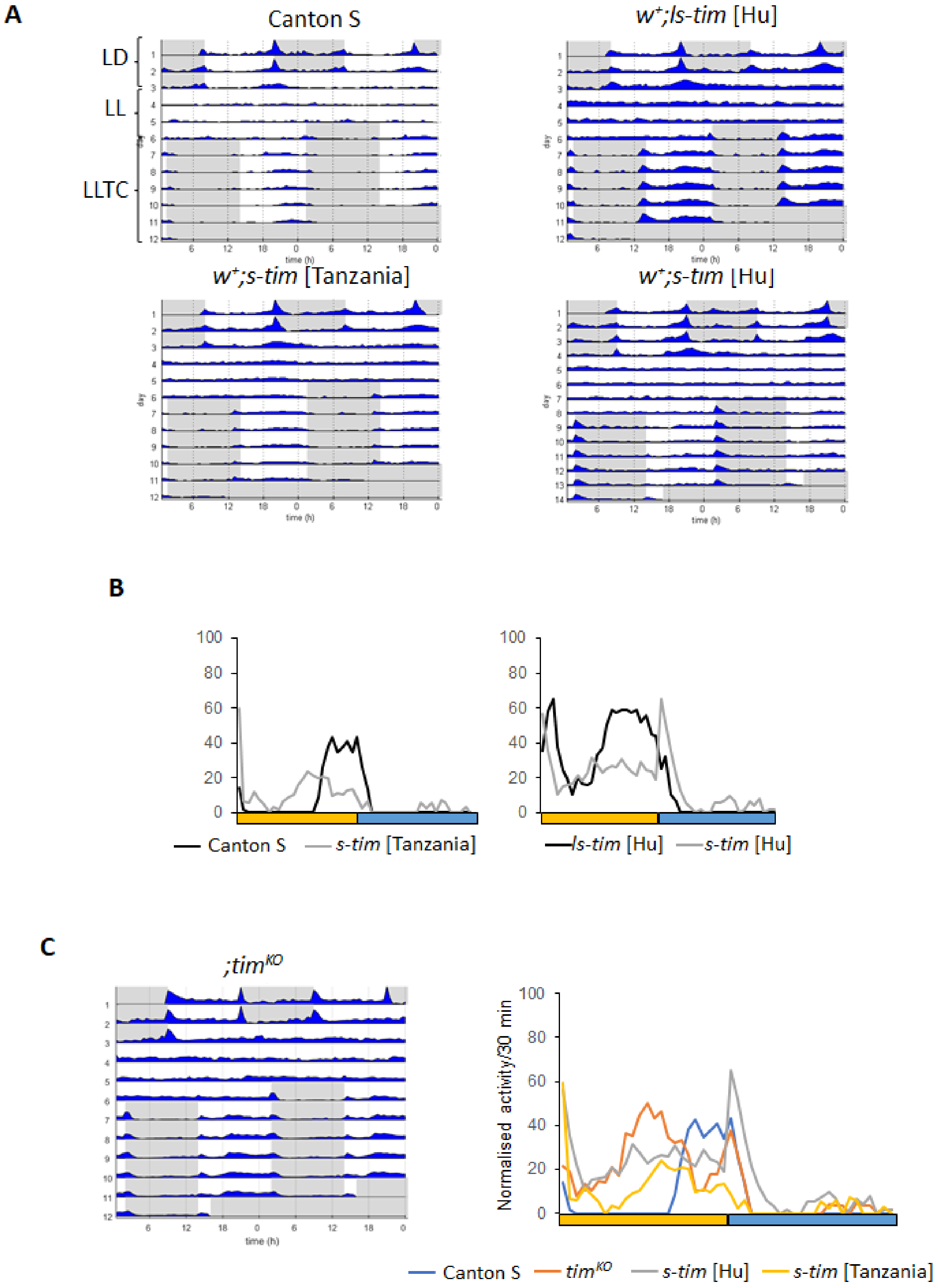

**Supplementary Figure 2: Eye colour influences the activity pattern of s-tim flies during constant light and temperature cycles. A)** Group actograms of one representative experiment as described in the legend for Figure 1A. N (Canton S): 20; (*$w^+$; ls-tim* [Hu]): 24; (*$w^+$; s-tim* [Tanzania]): 19; (*$w^+$; s-tim* [Hu]): 24. **B)** Median of normalised activity during day 6 of LLTC of independent experiments combined. Yellow bar: thermophase, blue bar cryophase (12h each). N (Canton S): 51; (*$w^+$; ls-tim* [Hu]): 43; (*$w^+$; s-tim* [Tanzania]): 19; (*$w^+$; s-tim* [Hu]): 63. **C)** Left: Group actogram of one representative experiment of $tim^{KO}$; Right: Median of the normalised activity. Same flies as in B with the addition of $tim^{KO}$ flies (N=56).

Supplementary Figure 3

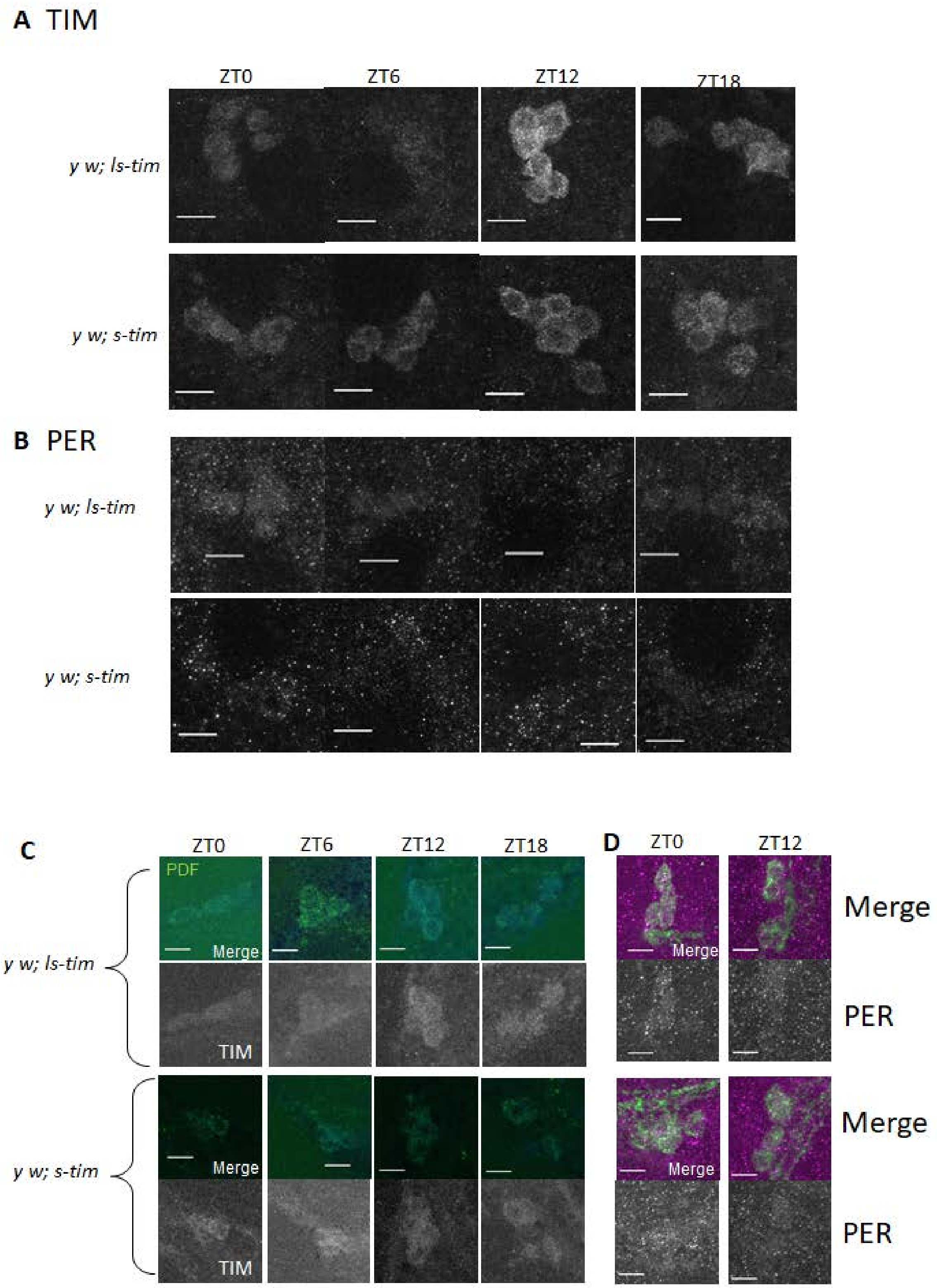


**Supplementary Figure 3: Constitutive cytoplasmic localisation of TIM and low PER levels in *s-tim* flies during constant light and temperature cycles. A-B)** TIM (A) and PER (B) in the

LNd on day six of LLTCin *y w; ls-tim* and *y w;s-tim* flies (for quantification see Figure 2). **C-D)** TIM (C, blue) and PER (D, magenta) in the s-LNv. PDF antibody (green) was used as a marker to identify the LN $PDF^{+}$ neurons. Scale bar: 10µm.

**Supplementary Figure 4**

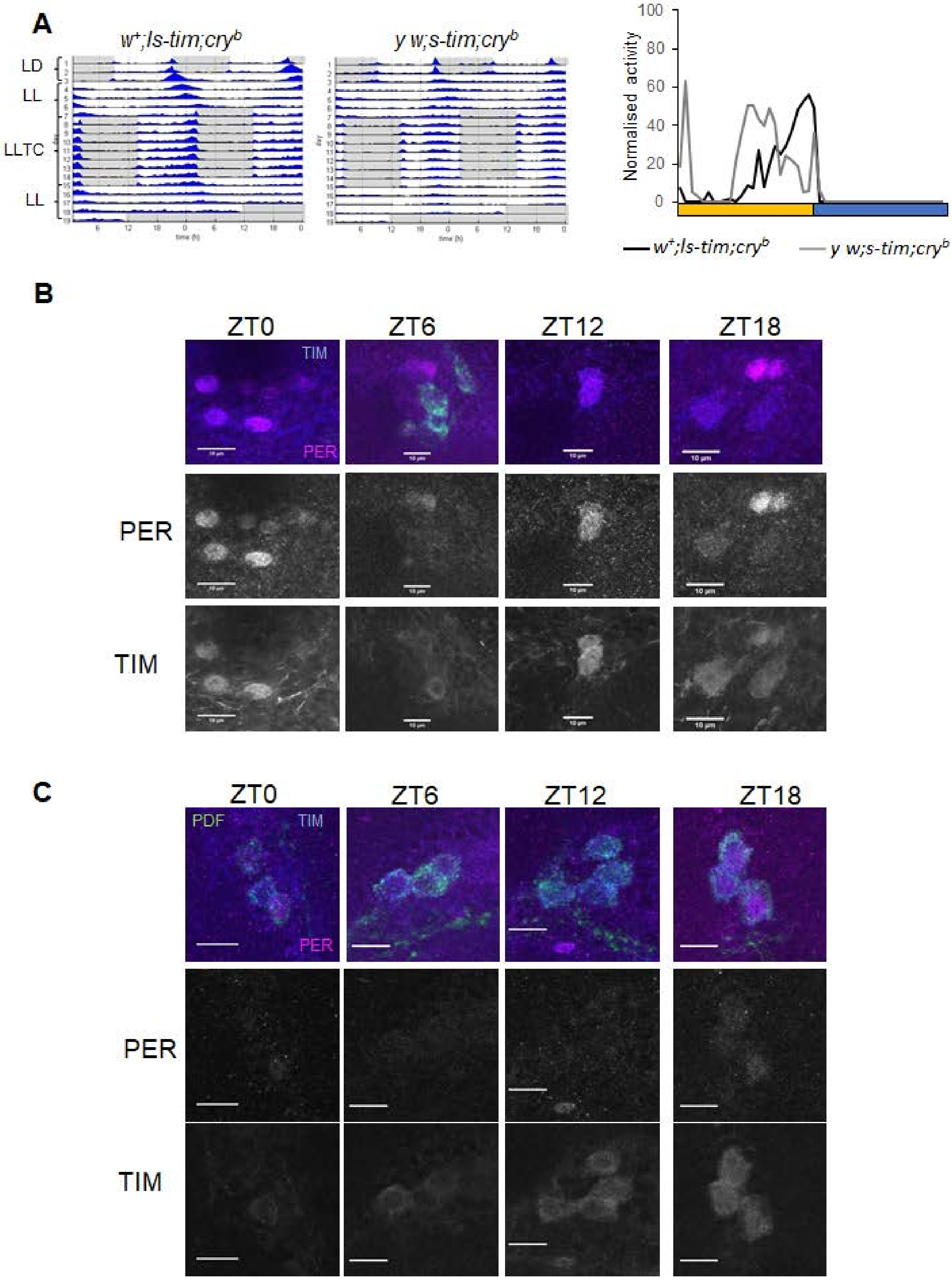


**Supplementary Figure 4: Cryptochrome depletion partially restores rhythmic behaviour molecular oscillations during constant light and temperature cycles in *s-tim* flies. A)** Group actograms (left) and median of the normalised activity of *w; ls-tim; cry$^b$* (N = 20) and *w; s-tim; cry$^b$* (N = 18) as described in the legend to Figure 1A, B, respectively. **B-C)** TIM (blue) and PER (magenta) in the LNd (B) and the s-LNv (C) on day six of LLTC. PDF antibody (green) was used as a marker to identify the LN PDF$^+$ neurons. Scale bar: 10μm.

**Supplementary Figure 5**

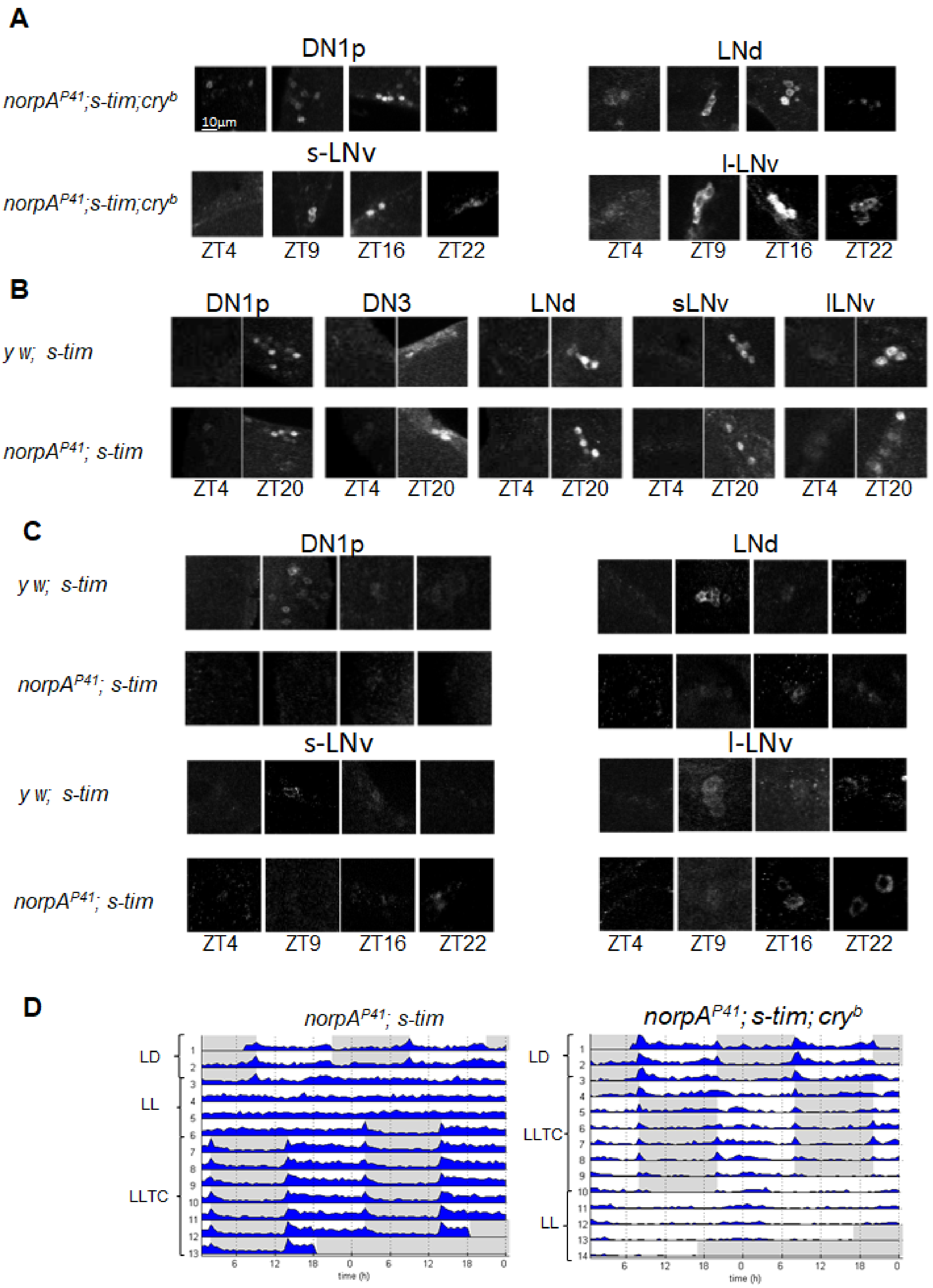

**Supplementary Figure 5: Visual system function and Cryptochrome prevent molecular and behavioural synchronisation of *s-tim* flies during constant light and temperature cycles. A)** Representative images of TIM staining at different time points during LLTC in *norpA*$^{P41}$*; s-tim ; cry*$^{b}$ flies. **B, C)** Representative images of TIM staining at different time points during DDTC (B) and LLTC (C) in *norpA*$^{P41}$*; s-tim* flies. **D)** Group actogram of *norpA*$^{P41}$*;s-tim* (N=20) and *norpA*$^{P41}$*; s-tim ;cry*$^{b}$ (N=15) flies as described in the legend to Figure 1A.

**Supplementary Figure 6**

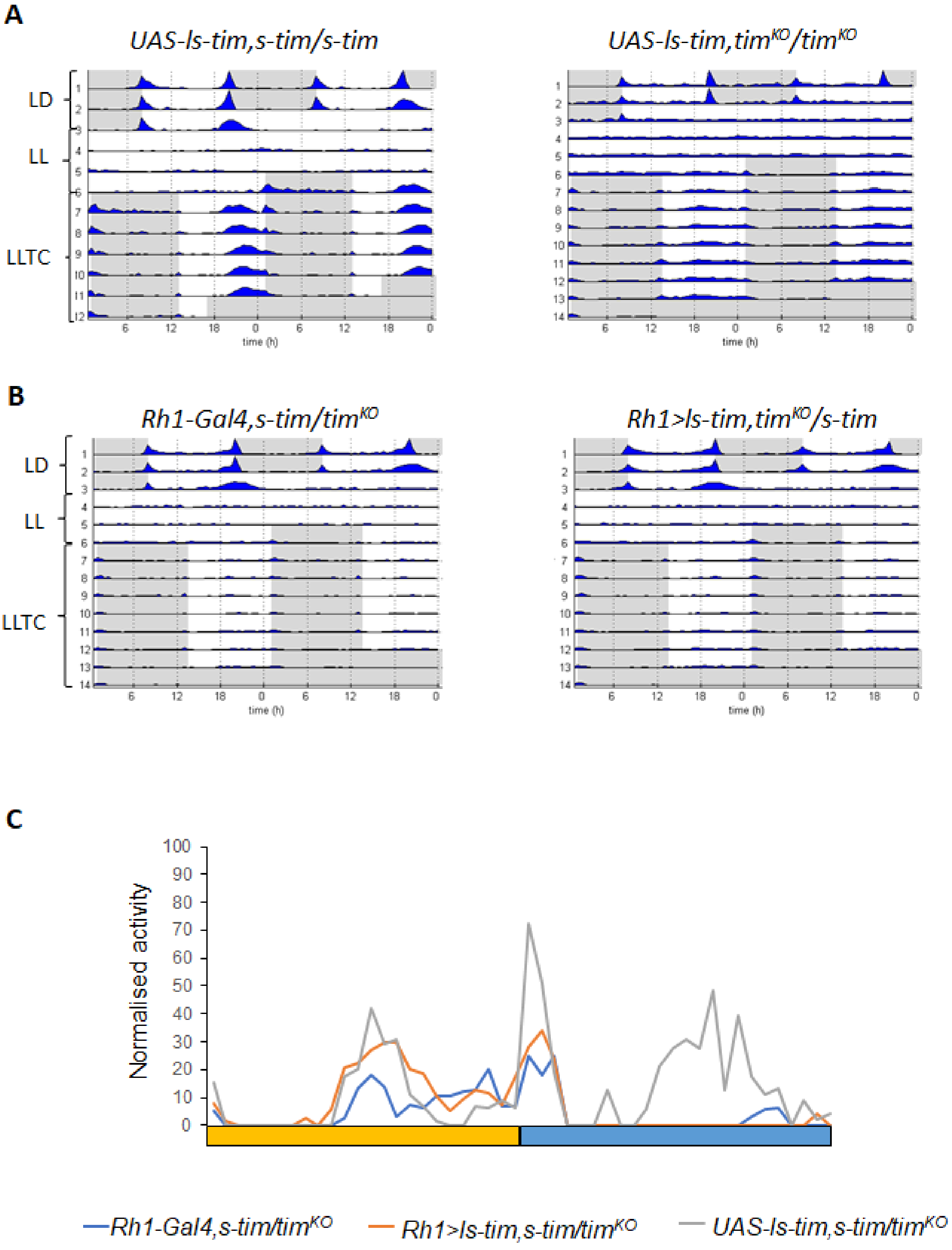


**Supplementary Figure 6: Expression of *ls-tim* in photoreceptor cells is not sufficient to restore synchronisation to temperature cycles during constant light in *s-tim* flies.** A) Group actograms of control flies carrying the *UAS-ls-tim* construct in a homozygous *s-tim* background (N = 20) and $tim^{KO}$ background (N = 17) as described in the legend to Figure 1A. Note that the *UAS-ls-tim* flies in homozygous *s-tim* background synchronize to LLTC (upper left), indicating leaky expression of *UAS-ls-tim* in absence of a *Gal4* driver. However, the level of expression is not sufficient to restore synchronisation to LLTC (and LD) in a homozygous $tim^{KO}$ background (upper right) or in heterozygous $tim^{KO}$/*s-tim* flies (Figure 5A). **B)** Group actograms as in (A) for *Rh1-Gal4, s-tim/*$tim^{KO}$ and *Rh1-Gal4, s-tim/UAS-ls-tim, s-tim,* $tim^{KO}$. N for both genotypes = 19. **C)** Median of normalised activity. Same flies as in B and Figure 5.

**Supplementary Figure 7**

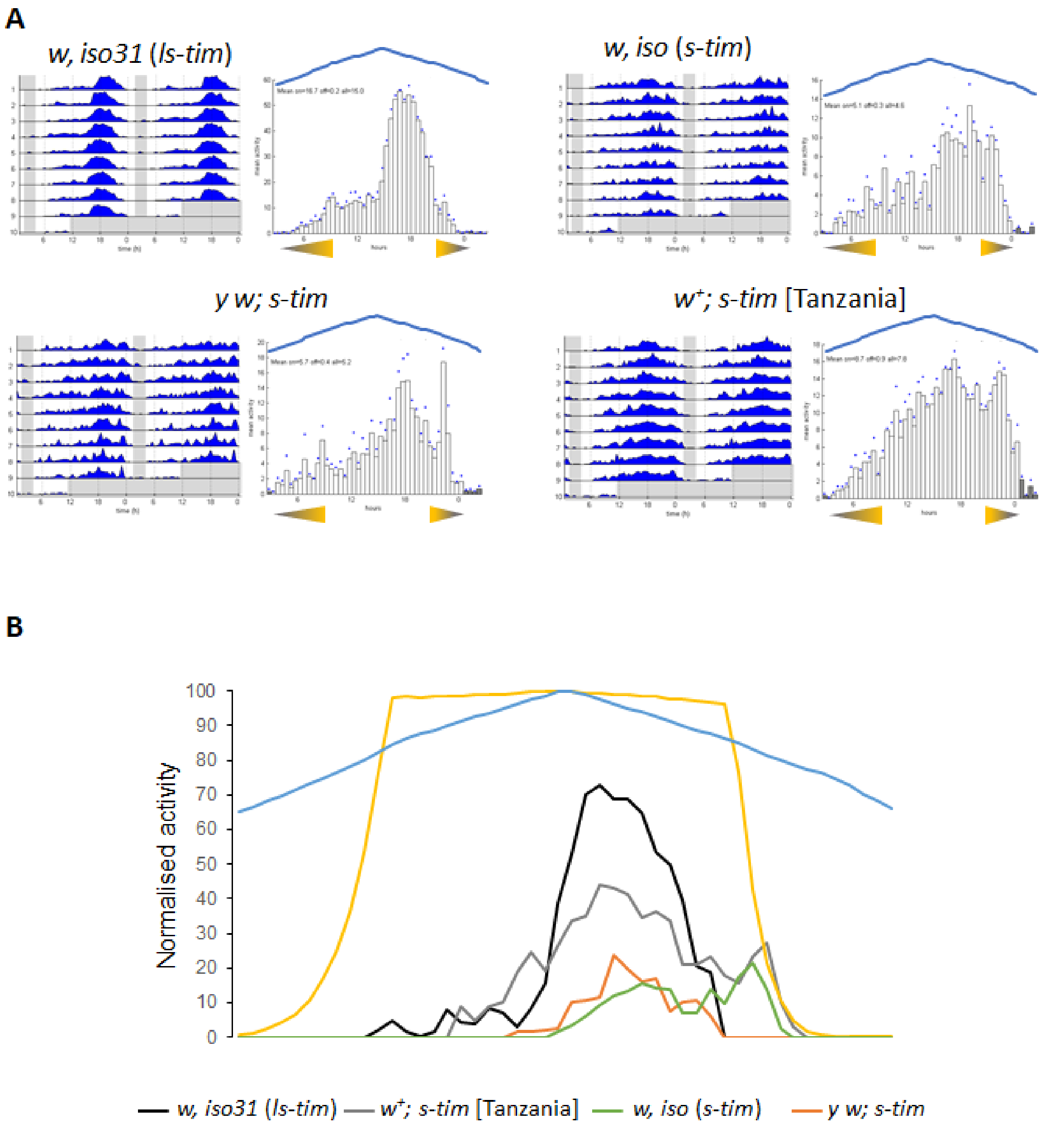


**Supplementary Figure 7: Only *ls-tim* flies are able to synchronize to Northern latitude summer conditions.** A) Group actograms and corresponding histograms of the last 3 days of one representative experiment as in legend to Figure 6B. N (*w, iso31 ls-tim*): 20; (*w, iso s-tim*): 20; (*y w; s-tim*): 19; (*w$^{+}$; s-tim* [Tanzania]): 16. **B)** Median of the normalised activity of two experiments on the 6$^{th}$ day under Oulu conditions. For all the genotypes, apart from *s-tim* [Tanzania] (N=16), N=40.
